# CAR T Cells Secreting IL18 Augment Antitumor Immunity and Increase T Cell Proliferation and Costimulation

**DOI:** 10.1101/111260

**Authors:** Biliang Hu, Jiangtao Ren, Yanping Luo, Brian Keith, Regina M. Young, John Scholler, Yangbing Zhao, Carl H. June

## Abstract

Interleukin 18 (IL18) is known to induce the expression of interferon-γ (IFNG), but its effects on T cell proliferation and costimulation are not completely understood. In this study, we demonstrate that ectopic expression of IL18 in CART cells caused significant T cell proliferation *in vitro* and *in vivo,* and enhanced antitumor effects in xenograft models. Moreover, IL18 mediated T cell expansion required neither tumor antigen nor CAR expression, and produced severe GVHD in NSG mice. Furthermore, recombinant IL18 costimulated IFNG secretion and proliferation of anti-CD3 beads treated T cells. Interestingly, IL18 costimulation could expand purified CD4 T cells, but not CD8 T cells. However, CD8 T cells proliferated greater than CD4 T cells in magnitude within bulk T cells, suggesting CD4 help effect was involved. Using CRISPR/Cas9 gene editing, we confirmed that IL18-driven expansion was both TCR and IL18 receptor (IL18R) dependent. Importantly, we demonstrated that TCR-deficient, IL18-expressing CD19 CART cells exhibited remarkable proliferation and persistent antitumor activity against CD19-expressing tumor cells *in vivo*, without eliciting any detectable GVHD symptom. Finally, we describe APACHE T cells, a novel strategy for coupling IL18 expression in CART cells to antigen stimulation, thereby limiting potential toxicity associated with persistent IL18 production. In sum, our study supports human IL18 as a T cell costimulatory cytokine for fueling CART therapy.

## Introduction

Following the initial reports of clinical success using chimeric antigen receptor (CAR) T cell therapy to treat B cell leukemia and lymphoma (Kochenderfer et al., 2010; Porter et al., 2011), there have been intense efforts to improve the design and clinical translation of CART cells (Dotti et al., 2014; Jensen and Riddell, 2014; Kalos and June, 2013). It is well established that T cell receptor (TCR) engagement and costimulatory signaling provide the critical signals that regulate T cell activation, proliferation and cytolytic functions (Chen and Flies, 2013). In addition to TCR and costimulatory signals, exogenous cytokines also play a fundamental role in modulating T cell function (Curtsinger and Mescher, 2010), as reflected in the influence of cytokines produced by CD4+ T helper cells on the activity of CD8+ cytotoxic T cells(Bevan, 2004). For example, interleukin 2 (IL2) promotes clonal expansion of effector T cells in response to antigen stimulation, and is also required to maintain suppressive regulatory T (Treg) cell-mediated immune tolerance (Boyman and Sprent, 2012).

CART cells are among the most promising of multiple adoptive T cell transfer-based immunotherapies currently under development (Harris and Kranz, 2016; Hinrichs and Rosenberg, 2014; Lu et al., 2014; Robbins et al., 2013). The manufacture of sufficient tumor-specific T cells in a clinical setting has relied on the use of exogenous cytokines, including IL2, IL7 andIL15, to promote *ex vivo* expansion of genetically engineered T cells (Cha et al., 2010; Cieri et al., 2013). In addition, IL21 has been shown to promote expansion and engraftment of adoptively transferred T cells (Chapuis et al., 2013).

Although many immune cytokines have been evaluated for their ability to promote anti-tumor immunity when delivered exogenously (Dranoff, 2004; Lee and Margolin, 2011), reports of toxicity and/or lack of efficacy have limited their success. Several attempts to replace exogenous cytokine treatment with T cell-autonomous cytokine production have been reported. For example, tethering IL15 to CART cells was shown to enhance T cell persistence and antitumor activity (Hurton et al., 2016); however, there are concerns that this approach could lead to transformation of the T cells (Hsu et al., 2007; Sato et al., 2011). Similarly, IL12-expressing T cells have been investigated in multiple preclinical studies (Chinnasamy et al., 2012; Chmielewski et al., 2014; Chmielewski et al., 2011; Koneru et al., 2015; Pegram et al., 2012; Zhang et al., 2015), and IL12-expressing tumor infiltrating lymphocytes (TIL) were recently used to treat melanoma patients (Zhang et al., 2015). Although IL12-expressing TIL therapy resulted in effective antitumor response, toxicity from persistent, elevated serum IL12 concentrations was observed. More recently, interferon-β (IFNB) has shown efficacy in bridging innate and adaptive immune responses (Yang et al., 2014), and may be an attractive option for engineering cytokine producing CART in future (Zhao et al., 2015).

We hypothesized that including an IL18 gene expression cassette in the CAR lentiviral vector might overcome the requirement for exogenous cytokine stimulation, and promote improved antitumor CART cell effects. IL18 was initially characterized as an inducer of interferon-γ (IFNG) expression in T cells (Nakamura et al., 1993; Nakamura et al., 1989),and was shown to activate lymphocytes and monocytes, without eliciting severe dose limiting toxicity, in a previous clinical study(Robertson et al., 2006). The combination of IL18 with IL12 treatment promotes Th1 and NK cell responses (Nakanishi et al., 2001), but also induces lethal toxicity in murine models (Carson et al., 2000; Osaki et al., 1998), reflecting the dynamic tension between achieving high potency and inducing toxicity in cytokine based immunotherapies. Despite initial indications that IL18 treatment alone may be only minimally effective in boosting anti-tumor immunity (Osaki et al., 1998), we previously reported that recombinant human IL18 could significantly enhance engraftment of human CD8 T cells in a xenograft model (Carroll et al., 2008).

In the current study, we demonstrate that T-cell autonomous IL18 expression, in combination with CAR or TCR signaling, promotes significant T cell proliferation *in vitro* and *in vivo.* Moreover, IL18 mediated T cell expansion in xenograft model required neither tumor antigen nor CAR expression, and produced severe GVHD in NSG mice. Interestingly, CD4+ T cells expanded preferentially in response to IL18 costimulation. Using CRISPR/Cas9 gene editing, we confirmed that IL18-dependent T cell expansion was both TCR and IL18 receptor (IL18R) dependent. Importantly, we demonstrated that TCR-deficient, IL18-expressing CD19 CART cells exhibited remarkable proliferation and persistent antitumor activity against CD19-expressing tumor cells and did not cause GVHD *in vivo*. These data support the redefinition of human IL18 as a T cell costimulatory cytokine and the utility of engineering CART cells to express IL18. We further describe an inducible expression system that couples IL18 expression levels to CAR stimulation, thereby reducing potential toxic effects of persistent IL18 expression in the absence of tumor antigen. Collectively, these studies identify a novel approach to enhancing CART cell expansion *in vivo,* which is a critical determinant of successful CART cancer immunotherapy.

## Results

### IL18 enhances proliferation of polyclonal antigen-specific and non-specific CAR T cells *in vivo*

To test the hypothesis that elevated IL18 signaling can enhance the anti-tumor effects of CART cell therapy, we introduced a gene expression cassette encoding human IL18 (or GFP as a negative control) into a previously described anti-mesothelin (SS1) CAR lentiviral construct (SS1-IL18) (Figure 1A). Similarly, the IL18 gene cassette was introduced into an anti-CD19 CAR construct (CD19-IL18) to serve as a control for antigen-specific effects (Figure 1A). SS1-IL18 CART cells efficiently expressed SS1 CAR and secreted active human IL18 into the culture medium, as assessed using an IL18 reporter cell line (Figure 1-figure supplement 1A,B,C).

**Figure 1.**
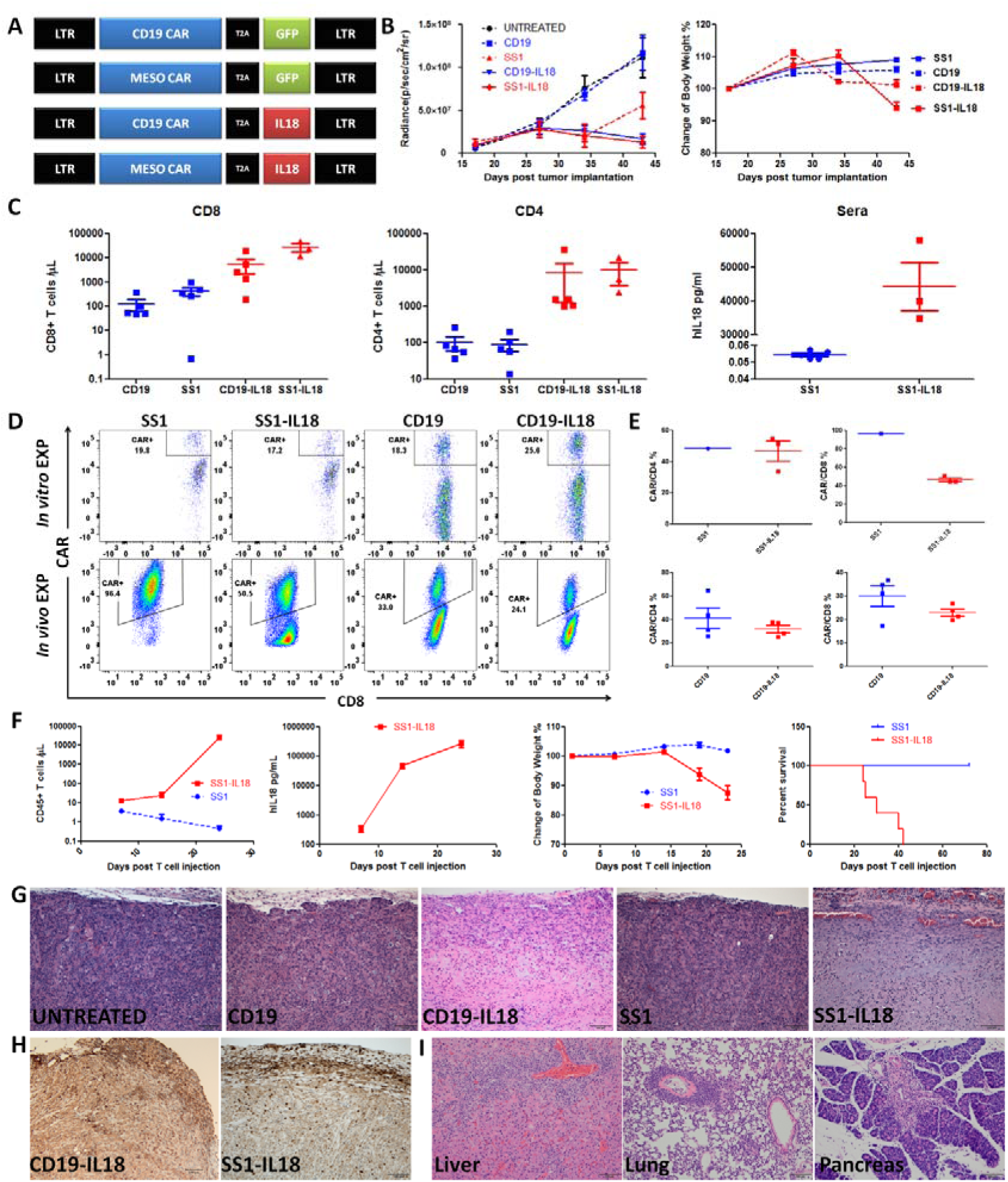
IL18 dependent T cell proliferation *in vivo,* and associated GVHD in NSG mice. **(A)** The construction designs of CART are listed as CD19-GFP, SS1-GFP, CD19-IL18, and SS1-IL18. **(B,C)** NSG mice (n=5) were inoculated with 2.5e6 AsPC1 expressing CBG-T2A-GFP. 3 weeks later, the mice received 2e6 of different CART as indicated. **(B)** Tumor growth and body weight. **(C)** CD8+ and CD4+ T cells in peripheral blood. IL18 levels were assessed by ELISA. **(D)** Representative FACS plots showing percentages of CAR+/CD8+ in CART cells during expansion *in vitro* and in the spleens of mice. Each data point represents cells from a single spleen, except that 5 spleens from mice receiving SS1 CART were pooled. **(E)** The percentage of CAR+/CD4+ or CAR+/CD8+ T cells in spleens from (D). **(F)** Tumor free NSG mice (n=5) were inoculated with 5e6 SS1 or SS1-IL18 CART cells, and monitored for circulating T cells, serum IL18 concentrations, body weight and survival. **(G)** H&E staining of tumors treated with different CART cells as indicated. **(H)** IHC staining for human CD3+ T cells in tumor sections from mice treated with CD19-IL18 or SS1-IL18 CART cells. **(I)** Representative pictures of liver, lung and pancreas from mice receiving SS1-IL18 CART cells demonstrated T cell infiltration and GVHD (see also Table S1). All data with error bars are presented as mean±SEM.

Interestingly, elevated IL18 expression enhanced the lytic activity of SS1-CART cells when cultured *in vitro* with the mesothelin-expressing BxPC3 pancreatic tumor cell line (Figure 1- figure supplement 1D). As previous data indicates that recombinant human IL18 significantly enhances engraftment of human CD8 T cells in xenograft models (Carroll et al., 2008), we investigated whether the IL18-expression could enhance CART cell-mediated antitumor effects *in vivo.* AsPC1 cells expressing click beetle green (CBG) and GFP reporter proteins were injected into NSG mice to generate xenograft tumors. Tumor-bearing mice were then injected with SS1, SS1-IL18, CD19 or CD19-IL18 CART cells, and tumor burden was monitored by bioluminescence. Surprisingly, both CD19-IL18 and SS1-IL18 CART cells suppressed tumor progression while inducing significant loss of body weight (Figure 1B), and CD19-IL18 or SS1-IL18 T cells were dramatically expanded 62-fold to 13993 cells/μl, or 74-fold to 38211 cells/μl, respectively (Figure 1C). Moreover, the serum IL18 concentration was increased to 44291pg/ml in SS1-IL18 CART cells (Figure 1C). These results indicate that CART cell-autonomous IL18 expression can drive expansion of both antigen-specific and non-specific CART cells in NSG mice.

To further analyze the specific effects of IL18 expression on CART cell proliferation, splenic T cells were analyzed for CAR expression by flowcytometry. Interestingly, in mice receiving SS1 CART cells, 96.4% of the *in vivo* expanded CD8 T cells were CAR+, whereas only 50% of the CD8+ T cells in mice receiving SS1-IL18 CART cells were CAR+ (Figure 1D,E). This expansion is more favorable to SS1 CAR+ T cells, as only approximately 24% of the expanded human T cells were CAR+ in mice receiving CD19-IL18 CART cells (Figure 1D,E). The percentage of CAR+ cells within CD4 or CD8 subsets are summarized in Figure 1E. These results suggested that IL18 promotes proliferation of both CAR+ and CAR- T cells in autocrine and paracrine modes. Phenotyping of TCRβ (TCRB) variants on expanded T cells reveals that all CART cell groups exhibit polyclonal expansion (Figure 1-figure supplement 2), suggesting that IL18 expression does not select for particular TCR specificities.

### IL18 mediated T cell proliferation induces lethal GVHD

Our observation that CD19-IL18 CART cells elicited antitumor activity and loss of body weight, as well as the expansion of untransduced T cells in mice receiving IL18-CART cells, suggested that IL18 expression in these contexts might promote severe GVHD, consistent with our previous report (Carroll et al., 2008). To test the severity of GVHD associated with IL18-CART cell therapy, we injected tumor-free NSG mice with either SS1 or SS1-IL18 CART cells, and monitored circulating human T cell levels, serum IL18 concentrations, body weight, and survival (Figure 1F). As SS1-IL18 CART cells expanded, mice exhibited significant loss of body weight, and started to die approximately 3 weeks after T cell injection (Figure 1F). Similarly, NSG mice bearing CD19- expressing Nalm6 tumors treated with CD19-IL18 CART showed dramatic expansion of human T cells in blood and spleen (Figure 1-figure supplement 3A,B). Remarkably, mice bearing AsPC1 cells succumbed quickly following injection with CD19-IL18 T cells, despite the absence of the CD19 tumor antigen on AsPC1 cells (Figure 1-figure supplement 3C). IL18 expression had no effect on the frequency of splenic regulatory T (Treg) cells (Figure 1-figure supplement 3D). Histological analyses confirmed that treatment with either SS1-IL18 and CD19-IL18 CART cells induced significant tumor destruction (Figure 1G). Immunohistochemical analysis for CD3 expression revealed the presence of T cells in the tumor treated with IL18-CART (Figure 1H), and necropsy of mice receiving SS1-IL18 CART cells (Table S1), revealed remarkable GVHD in liver and lung and atypical lymphocyte infiltration in pancreas (Figure 1I). Collectively, our results suggest that expressing IL18 in the context of CART cell therapy can enhance antitumor effects and xenogeneic GVHD, but may also produce unwanted toxicity due to TCR-mediated autoimmune effects in an autologous setting, or GVHD in an allogeneic setting. To optimize the use of IL18 in CART cell therapy, it was therefore important to dissect the contribution of TCR signaling to IL18-mediated CART cell effects *in vitro* and *in vivo.*

### TCR activation and IL18 promotes CD4 helper dependent T cell expansion *in vitro* and *in vivo*

Based on our observation that IL18 expressing T cells proliferated in NSG mice independent of CAR activation, and produced xenogeneic GVHD, we speculated that a functional synergy exists between IL18 and TCR stimulation. To test this hypothesis *in vitro,* primary human T cells were stimulated using anti-CD3 beads and recombinant IL18. Whereas combined exposure to anti-CD3 beads and IL18 efficiently stimulated increases in T cell size and proliferation (Figure 2A,B), neither treatment alone was effective, although anti-CD3 beads alone caused intermediate increase in size without triggering T cell proliferation. These results were confirmed in T cells isolated from two normal donors (Figure 2C).

**Figure 2.**
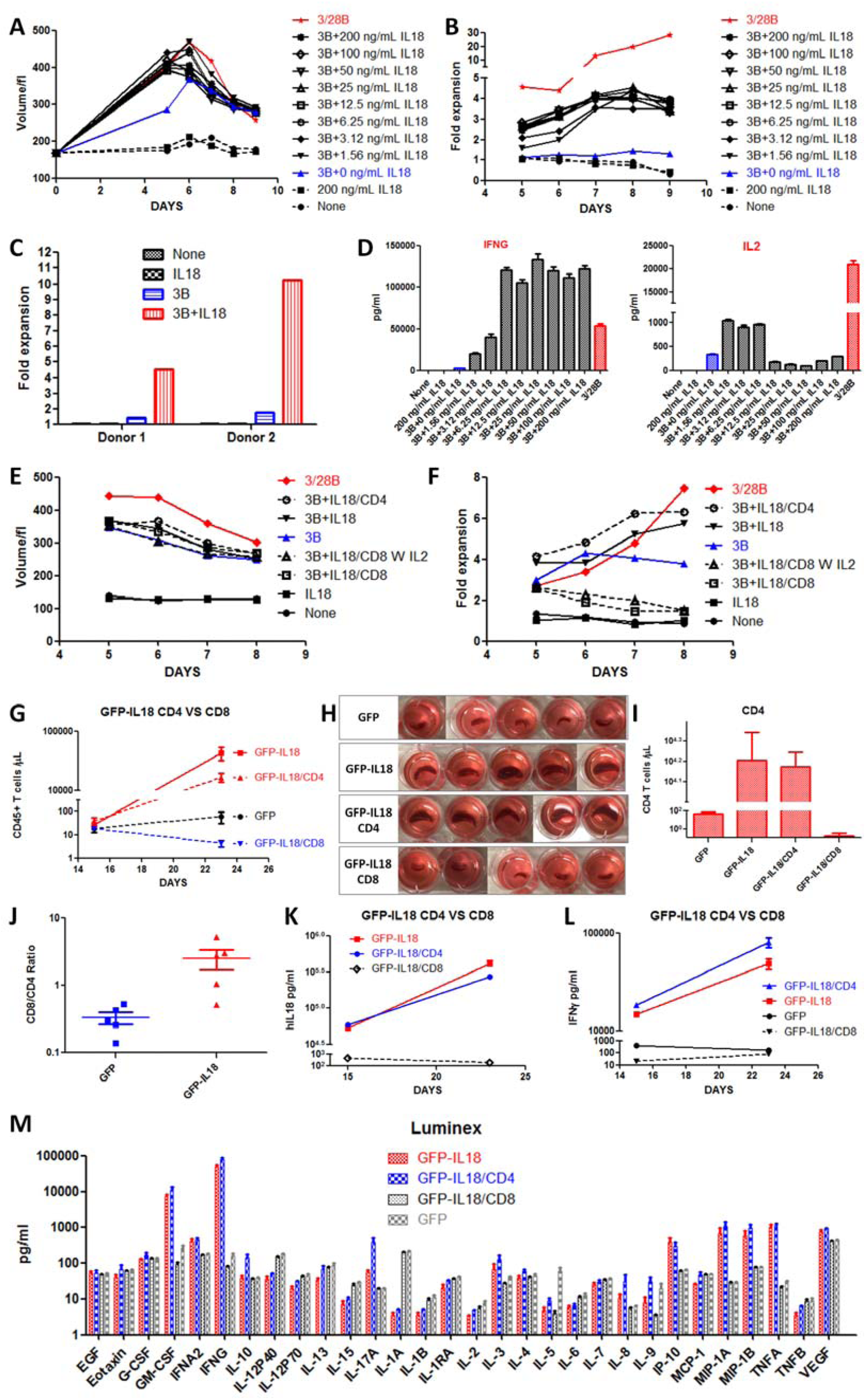
Combined TCR and IL18 signaling promotes CD4 helper dependent T cell expansion. (A-D) Primary T cells from normal donors were activated with anti-CD3 beads and varying concentrations (1.56 doubled up to 200 ng/ml) of recombinant IL18 (added at D0 and D3). Activated T cells were de-beaded at D5 and counted every day afterwards. **(A)** changes in T cell size and **(B)** proliferation kinetics. **(C)** IL18 dependent expansion of T cells from two different donors. **(D)** Luminex analysis of secreted IFNG and IL2. Additional cytokine data are shown in Figure 2-figure supplement 1. **(E,F)** Normal donor T cells expanded with anti-CD3/CD28 beads were selected by MACS columns to purify CD8 and CD4 T cells. Purified CD8, CD4 T cells, or bulk CD8+CD4 T cells were restimulated with anti-CD3 beads and 100ng/ml IL18 (added at D0 and D3). Cell size **(E)** and proliferation kinetics **(F)** were monitored after stimulation with anti-CD3 beads and recombinant IL18. **(G-M)** T cells were transduced with lentivector encoding GFP or GFP-IL18 at D1 during standard T cells expansion and GFP-IL18/CD8 and GFP-IL18/CD4 T cells were selected using MACS columns. NSG mice (n=5) were injected intravenously with 2×10^6^ GFP-IL18 T cells, GFP-IL18/CD8, or GFP-IL18/CD4 T cells. GFP T cells were included as a negative control. The data shown is representative of two independent experiments. **(G)** Circulating CD45+ T cells at d15 and D23 after T cells injection. **(H)** Spleens from various groups, indicating splenomegaly in mice receiving GFP-IL18/CD4 or bulk T cells. **(I)** Circulating CD4+CD45+ T cells at D23 following T cell injection. **(J)** The ratio of CD8 to CD4 T cells. Circulating Serum levels of IL18 **(K)** and IFNG **(L)** at D15 and D23. **(M)** Serum at D23 was analyzed by luminex (duplicate) to assess cytokine profile.

To better characterize IL18 activation of T cells, we collected the culture supernatants at D5 from different conditions as indicated to analyze the profile of secreted cytokines. As expected, low concentrations of IL18 combined with anti-CD3 beads, significantly enhanced IFNG production compared to beads alone (Figure 2D). Although low concentrations of IL18 and anti-CD3 beads treatment enhanced IL2 production, maximal IL2 levels were at least 20-fold lower than those induced by standard anti-CD3/28 beads (Figure 2D). The production of other cytokines was generally comparable between CD3/IL18 and CD3/D28 activation (Figure 2-figure supplement 1). Multicolor flow phenotype revealed that IL18-expanded T cells had a primarily central memory (CM) phenotype (CCR7+CD45RO+), similar to that obtained with anti-CD3/28 beads (Figure 2-figure supplement 2).

To address the specific roles of T cell subsets in these responses, T cells expanded by standard anti-CD3/CD28 beads were sorted by magnetic-activated cell sorting (MACS) to generate CD4 and CD8 subpopulations. CD8 T cells failed to proliferate after restimulation with IL18 and anti-CD3 beads despite a significant increase in size, and addition of IL2 had no additional effect (Figure 2E,F). In sharp contrast, CD4 T cells or bulk T cells restimulated with IL18 and anti-CD3 beads proliferated efficiently, at a level comparable to that of combined CD4+CD8 T cells following CD3/CD28 restimulation. Restimulated CD4 and CD8 T cells displayed transiently elevated central memory (CM) markers, but ultimately expressed a primarily effector memory (EM) phenotype (Figure 2-figure supplement 3A), similar to that observed for restimulated IL18- CART cells (Figure 2-figure supplement 3B).

GFP-IL18 expressing CD8 T cells also failed to expand *in vivo* following injection into NSG mice (Figure 2G). In contrast, GFP-IL18 expressing CD4 or bulk T cells exhibited greater than 400-fold expansion between days 15 to 23 post T cell injection (Figure 2G), and were associated with enlarged spleens (Figure 2H). The number of expanded CD4 T cells was comparable whether pure CD4 T cells or total T cells were injected, suggesting that CD8 T cells do not have a significant effect on CD4 T cell expansion (Figure 2I). Interestingly, CD8 T cells proliferated greater than CD4 T cells in magnitude within bulk T cells, suggesting CD4 help effect was involved (Figure 2J).

Serum cytokine levels were determined in mice receiving CD4, CD8, or bulk T cells expressing GFP-IL18, or control GFP T cells. Consistent with our *in vitro* results, serum IL18 levels increased in the GFP-IL18 T cell and GFP-IL18 CD4 T cell populations, but not in the isolated GFP-IL18 CD8 T cell population (Figure 2K). IFNG levels also increased in proportion to T cell expansion (Figure 2L), and GM-CSF levels were significantly elevated in the expanding GFP-IL18 T cell and GFP-IL18 CD4 T cell populations, whereas other cytokines failed to show large changes (Figure 2M). In summary, these results suggested that IL18 acts as a costimulatory cytokine that interacts synergistically with TCR activation to drive efficient T cell proliferation, largely dependent on effects through CD4 helper T cells.

### CAR or TCR signaling is required for IL18 mediated T cell proliferation *in vitro* or *in vivo*

To first evaluate the effect of IL18 coexpression on CART cell proliferation *in vitro,* we deleted TCR mediated allogeneic effect in SS1-IL18 and CD19-IL18 CART cells using CRISPR/Cas9 gene editing as previously described (Ren et al., 2016). TCR KO IL18-CART cells had only marginally enhanced expansion following CD3/CD28 beads activation and RNA electroporation (Figure 3A). When stimulated with irradiated K562 cells expressing cognate antigen and exogenous IL2, SS1-IL18 CART cells displayed significantly increased proliferation relative to SS1 CART cells (Figure 3B), which were producing high level of IFNG and IL2, and polyclonal in TCRB variants distribution (Figure 3-figure supplement 1A,B,C). However, CD19-IL18 CART cells expanded slightly better than the CD19 CART cell control (Figure 3C). Interestingly, IL18 expression promoted a modest increase in proliferation in both TCR-deficient SS1-IL18 and CD19-IL18 CART cells relative to their TCR wild type counterparts (Figure 3B,C,D). Robust IL18 secretion was observed in all cases (Figure 3E, Figure 3-figure supplement 2A,B), while IL18-CART displayed enhanced cytokine production of IFNG and several others (Figure 3-figure supplement 2C). The observation of extended proliferation in SS1-IL18 CART should be attributed to antigen independent activation of SS1 CAR as previously found (Frigault et al., 2015). Moreover, the loss of proliferation in TCR-deficient SS1-IL18 CART cells suggested that TCR structure might be necessary to maintain the antigen independent activation of SS1 CAR. Nevertheless, this data strongly suggested that IL18 could drive persistent T cell proliferation in presence of continuous CAR activation *in vitro.*

**Figure 3.**
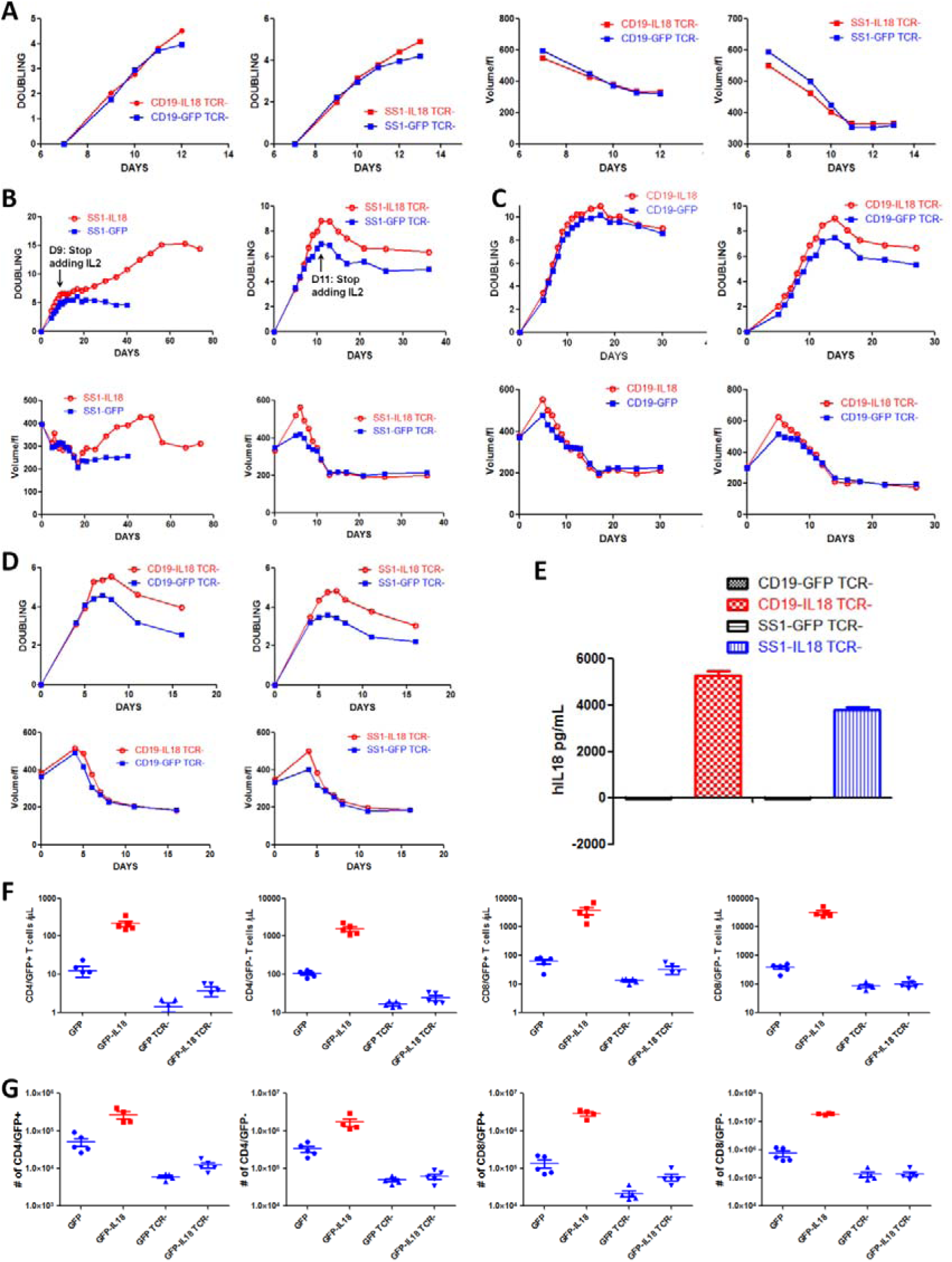
IL18 mediated T cell proliferation requires CAR or TCR signaling. **(A)** Expansion kinetics and size of TCR-deficient (TCR-) CD19-GFP or CD19-IL18 CART cells following stimulation with anti-CD3/CD28 beads and RNA electroporation. **(B)** Expansion kinetics and size of TCR+ and TCR-deficient SS1-GFP or SS1-IL18 CART cells following restimulation with irradiated mesothelin-expressing K562 cells in the presence of exogenous IL2 (until day 9 or day 11, as indicated). **(C)** Expansion kinetics and size of TCR+ and TCR-deficient CD19-GFP or CD19-IL18 CART cells following restimulation with irradiated CD19-expressing K562 cells in the presence of exogenous IL2. **(D)** TCR-deficient CART cells expanded from (B) and (C) were further restimulated with aAPC in the absence of exogenous IL2. **(E)** IL18 secretion (duplicate) of indicated CART cells following restimulation with cognate antigen. **(F,G)** NSG mice (n=5) were injected with 2×10^6^ TCR+ or TCR-deficient T cells expressing either GFP or GFP-IL18. After 18 days, the number of GFP+ and GFP-CD4+ and CD8+ cells in blood **(F)** or spleens **(G)** was assessed. All data with error bars are presented as mean±SEM.

To test whether continuous TCR activation by mouse MHC molecules was required for general IL18-mediated T cell expansion *in vivo,* NSG mice were injected with TCR+ and TCR-GFP-IL18 T cells. TCR+ GFP-IL18 expressing T cells expanded dramatically compared to TCR+ GFP control T cells, whereas TCR-GFP-IL18 expressing T cells showed negligible IL18-induced expansion relative to controls (Figure 3F,G). In addition, flowcytometry analysis of spleen cells indicated that TCR+ GFP-IL18 expressing T cells displayed high levels of the IL18 receptor (IL18R) (Figure 3- figure supplement 3A), and displayed a primarily effector memory phenotype (Figure 3-figure supplement 3B). Collectively these data indicate that CAR or TCR activation is required for IL18- mediated T cell proliferation *in vitro* or *in vivo,* respectively.

### IL18 receptor is required for cell-autonomous IL18-medated T cell expansion *in vivo*

To further investigate the synergy between IL18 and TCR signaling, we generated IL18Rα (IL18RA) deficient T cells using CRISPR/Cas9 mediated gene editing (Figure 4-figure supplement 1A,B). We next employed a cell competition assay using four distinct populations of engineered T cells (Figure 4A). These included (1) WT CD19-IL18 T cells; (2) WT GFP T cells; (3) TCR KO T cells labeled with DsRed; and (4) IL18R KO T cells labeled with AmCyan. Successful deletion of the TCR and IL18R was confirmed by flowcytometry, and MACS depletion was performed to remove any residual T cells expressing either TCR or IL18R proteins (Figure 4B). Following *ex vivo* expansion, all four T cell populations were mixed and flowcytometry analysis confirmed the ratio of (WT CD19-IL18):(WT GFP):(TCR KO DSRED):(IL18R KO AMCYAN) cells to be 8.59% : 7.45% : 6.71% : 7.27% of total T cells (Figure 4C). The mixed population of T cells was injected into NSG mice and blood and spleen were collected for analysis when mice exhibited significant loss of bodyweight. Staining for IL18RA indicated that AmCyan labeled T cells partially regained low levels of IL18R expression (Figure 4D), and likely represent T cells that did not undergo successful gene editing and were not completely removed by MACS depletion.

**Figure 4.**
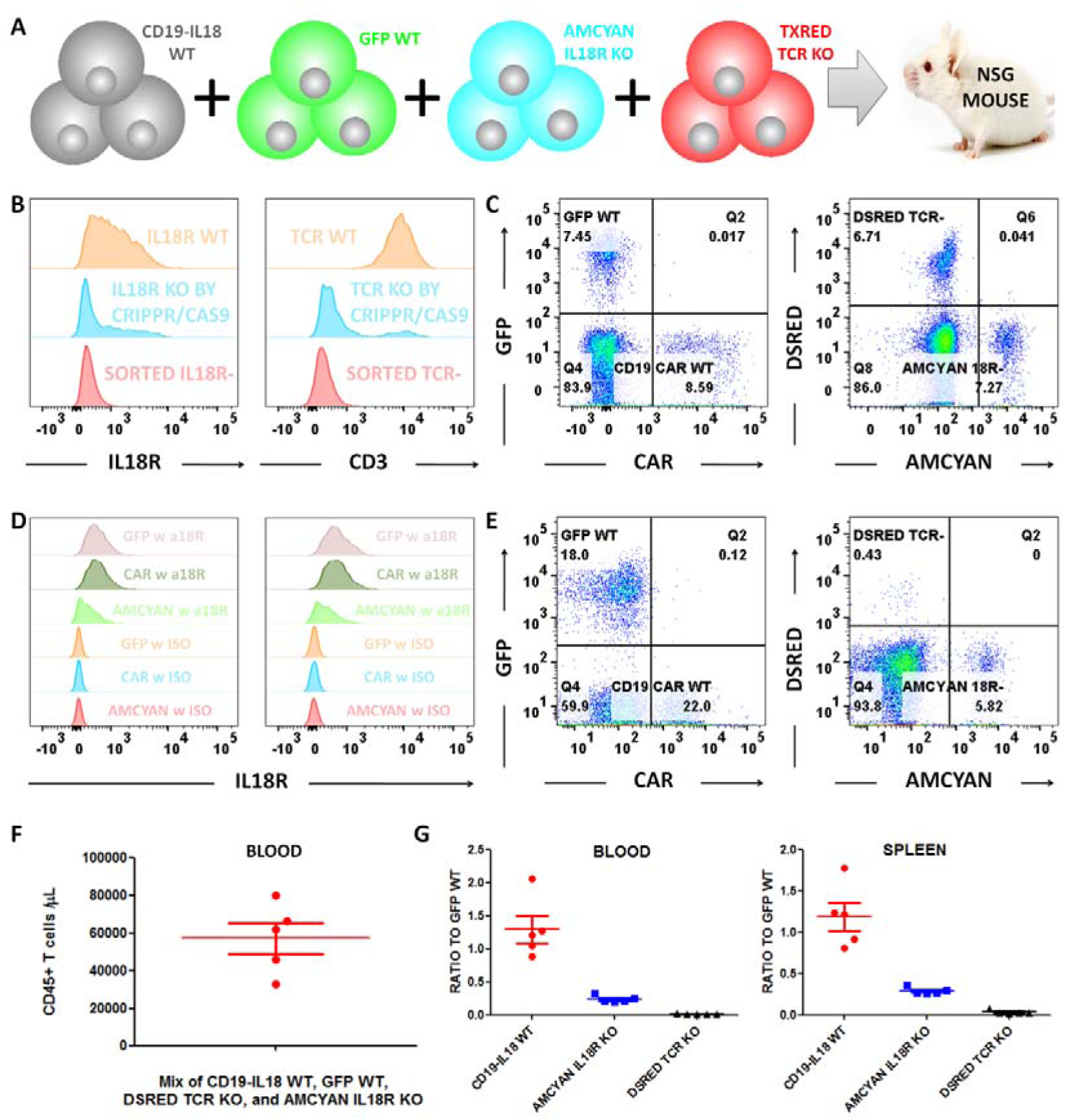
IL18 receptor is required for cell-autonomous IL18-mediated T cell expansion *in vivo.* **(A)** Cell competition assay design. CD19-IL18 WT, GFP WT, Amcyan IL18R KO, and DsRed TCR KO T cells were mixed together and injected into NSG mouse (n=5) intravenously, to evaluate the expansion of each cell population in response to IL18 secretion. **(B)** Normal human primary T cells were activated and transduced with lentiviral expression vectors by standard protocols, then electroporated with sgRNA targeting TCRB or IL18RA, and Cas9 RNA as described in Methods. Three days after electroporation, T cells were stained with anti-CD3 or anti-IL18RA to confirm gene knockout. T cells with TCR or IL18R knockout were stained with anti-CD3, or anti-IL18RA microbeads and negatively sorted by MACS. **(C)** 1×10^6^CD19-IL18 CART cells, 1×10^6^GFP T cells, 1×10^6^IL18R deficient Amcyan T cells, and 1×10^6^TCR-deficient DsRed T cells were mixed together and injected intravenously into NSG mice. **(D)** Expression levels of IL18RA on GFP WT, CD19 CAR WT, and Amcyan IL18R- cells in blood (left) and spleen (right). **(E)** Frequencies of CD19-IL18 WT, GFP WT, IL18R-deficient Amcyan, and TCR-deficient DsRed T cells in blood. **(F)** Total circulating CD45+ cells, indicating robust expansion. **(G)** The ratios of CD19 CART WT, IL18R-deficient Amcyan, and TCR-deficient DsRed to GFP T cells in blood and spleen. All data with error bars are presented as mean±SEM.

Flowcytomety analysis revealed the final ratio of (WT CD19-IL18):(WT GFP):(TCR KO DSRED):(IL18R KO AMCYAN) to be 22.0% : 18.0% : 5.82% : 0.43% (Figure 4E). The number of CD45+ human T cells in individual mice ranged from 32800 to 79600 cells/μl blood (Figure 4F), confirming that all mice exhibited IL18-mediated T cell proliferation. Interestingly, the ratio of WT CD19-IL18 to WT GFP T cells following *in vivo* expansion was essentially unchanged (Figure 4G), consistent with the data in Figure 1D,E indicating that IL18 could drive equal expansion of CD19 CAR positive and negative T cells. In contrast, the proportion of IL18R KO AMCYAN T cells decreased significantly in blood and spleen, and it was even more dramatic for TCR KO DSRED T cells (Figure 4G). These data indicate that both TCR and IL18R signaling are required for IL18 mediated proliferation of human T cells as tested *in vivo* in the NSG model.

### IL18 costimulation with CAR signaling significantly enhanced CART proliferation and antitumor activity in xenograft model

As our data indicate that IL18 expression promotes TCR-mediated T cell proliferation, we hypothesized that IL18 costimulation would also enhance CART cell proliferation and anti-tumor efficacy. To avoid GVHD, only TCR-deficient (TCR-) T cells were used in these experiments. NSG mice were engrafted with Nalm6 tumor cells expressing CBG and GFP reporter proteins, and subsequently treated with either TCR- CD19-IL18 or control TCR- CD19-GFP CART cells. Tumor burden was assessed by bioluminescent imaging. TCR- CD19-IL18 T cells effectively cleared detectable tumor cells by day 10 (Figure 5A), in contrast to TCR- CD19-GFP T cells (Figure 5A,B). In addition, TCR-SS1-IL18 CART cells had no effect on tumor progression (Figure 5B). Although TCR- CD19-IL18 CART cells caused a transient decrease in body weight at D8, body weight returned to baseline levels and remained stable thereafter (Figure 5C). Remarkably, mice injected with TCR- CD19-IL18 CART cells remained viable for over 185 days post T cell injection, except for one mouse that exhibited tumor relapse at day 101 (Figure 5A,D) due to downregulation of CD19 expression (Figure 5-figure supplement 1A). Emergence of contaminating TCR+ CD19-IL18 CART cells was not observed at any point during treatment (Figure 5G).

**Figure 5.**
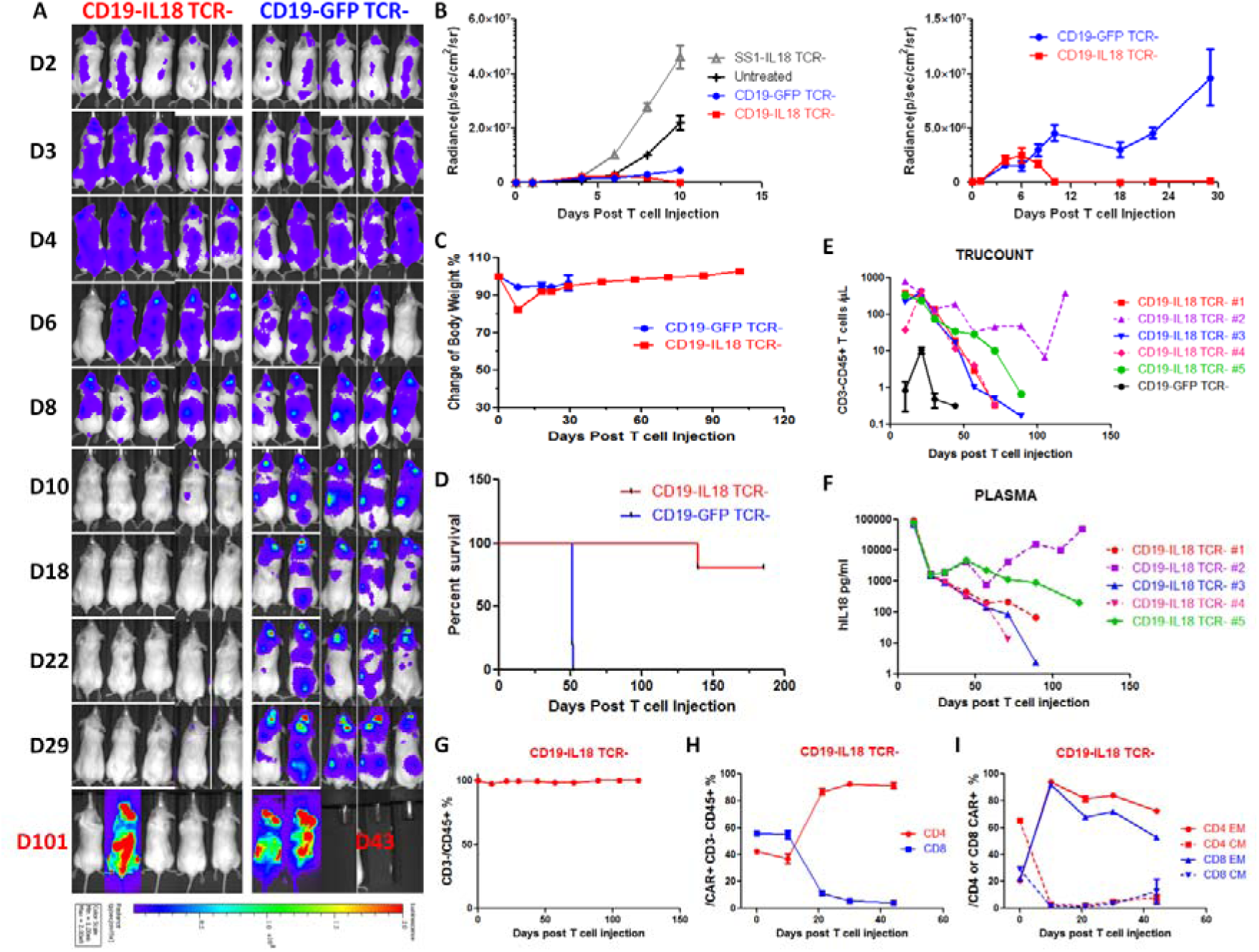
IL18 costimulation significantly enhanced proliferation and antitumor tumor activity of CD19 CART cells. NSG mice were injected intravenously with 1×10^6^ Nalm6 cells expressing CBG and GFP reporter proteins. At D7 post tumor injection, mice (n=5) were imaged and randomized into groups to receive 1×10^6^TCR-deficient CD19-GFP T cells or CD19-IL18 CART cells. Mice receiving no CART cells or TCR-deficient SS1-IL18 CART cells were included as controls. The data shown is representative of two independent experiments. **(A)** Live animal imaging of CBG expressing tumors in mice receiving CD19-GFP or CD19-IL18 TCR-deficient CART cells. **(B)** Left panel: tumor growth curves for mice injected with CD19-GFP CART cells, SS1-IL18 CART cells, or CD19-IL18 CART cells (all TCR-deficient). Untreated control received no T cells. Right panel: tumor growth in mice receiving TCR-deficient CD19-GFP CART cells or TCR-deficient CD19-IL18 CART cells. Mice were monitored for body weight **(C)** and survival **(D)**. The number of CD3-CD45+ T cells **(E)** and IL18 concentration **(F)** in blood were monitored over time. **(G)** The percentage of CD3-cells among total CD45+ cells, distribution of CD4+ and CD8+ T cells within CD19 CART cells **(H)**, and memory phenotype in CD4+ and CD8+ CART cells **(I)** in mice receiving TCR-deficient CD19-IL18 CART cells. All data with error bars are presented as mean±SEM.

From day 10 through day 21, mice receiving TCR- CD19-IL18 CART cells maintained approximately 350 CD3-CD45+ T cells/μl blood (Figure 5E), whereas the number of CD19-GFP CART cells varied between0.8 and 10.5 CD3-CD45+ T cells/μl. This approximately 100-fold difference suggests that IL18 significantly enhanced of proliferation CD19 CART cells, resulting in superior antitumor activity. Consistent with the expansion of CD19 CART cells, IL18 concentrations were high, averaging 77500pg/ml at D10 (Figure 5F). Subsequently, the number of T cells and IL18 concentrations decreased to below 1 CD3-CD45+ T cell/μl and approximately 100pg/ml in tumor-free mice. Flowcytometric analyses revealed that the distribution of CD4 and CD8 T cells rapidly evolved from equal balance at D10 (37% vs 55%, respectively) to CD4 domination at D21 (86% vs 11%) (Figure 5H). Interestingly, CD19-IL18 CART cells attained a primarily EM phenotype in both CD4 and CD8 subsets (Figure 5I). In the single mouse that developed a recurrent tumor associated with CD19 downregulation (Figure 5-figure supplement 1A), the number of T cells rose back to 373 CD3-CD45+ T cells/μl, and IL18 concentration back to 50209 pg/ml, while the percentage of CD3-CD45+ cells was still 99.6% (Figure 5-figure supplement 1B). This observation suggested that TCR- CD19-IL18 CART cells were still capable of expanding in response to tumor relapse, presumably in an antigen-dependent manner, while retaining a primarily CD4 T_EM_ phenotype (Figure 5-figure supplement 1C).

These results collectively demonstrate that IL18 can significantly enhance CART cell proliferation and induce superior anti-tumor activity. Equally important, the use of TCR-deficient CART cells obviated concerns of developing TCR-mediated GVHD during therapy.

### Designing an NFAT-based inducible system for IL18 production: APACHE T cells

Our data indicate that including the IL18 expression cassette within the CAR lentiviral design strongly enhanced T cell proliferation in response to TCR or CAR stimulation, and may be preferable to expanding CART cells that do not constitutively secrete cytokines. However, the high concentration of systemic IL18 produced by CART cells could raise potential safety concerns related to pathological activation or proliferation of bystander T cells reacting against foreign or self-antigens, or induction of NK cell mediated inflammation. We therefore designed an inducible system in which IL18 production is only activated after CAR engagement with tumor antigen. Specifically, we generated a lentiviral vector in which the expression of CD19 CAR, GFP, and IL18 is under the control of the NFAT promoter (Figure 6A). Once expressed, CD19 CAR will engage with antigen to stimulate NFAT promoter activity. This, in turn, will maintain CD19 CAR and IL18 expression through a positive feedback circuit. Once tumor antigen is cleared, CD19 CAR and IL18 expression levels should return to baseline, overcoming potential toxic effects of long-term constitutive IL18 expression. Because the endogenous TCR can also activate NFAT signaling pathways, we conducted these experiments in T cells rendered deficient in TCR expression by CRISPR/Cas9. In order to initially activate the signaling cascade, the cells were engineered to transiently express a CD19 CAR by mRNA electroporation into TCR-deficient T cells already lentivirally transduced with the NFAT-driven cassette (Ren et al., 2016; Zhao et al., 2010), as shown in Figure 6A. Deleting the TCR gene in these cells by CRISPR/Cas9 gene editing avoids unwanted NFAT activation by endogenous TCR signaling (Figure 6A).

**Figure 6.**
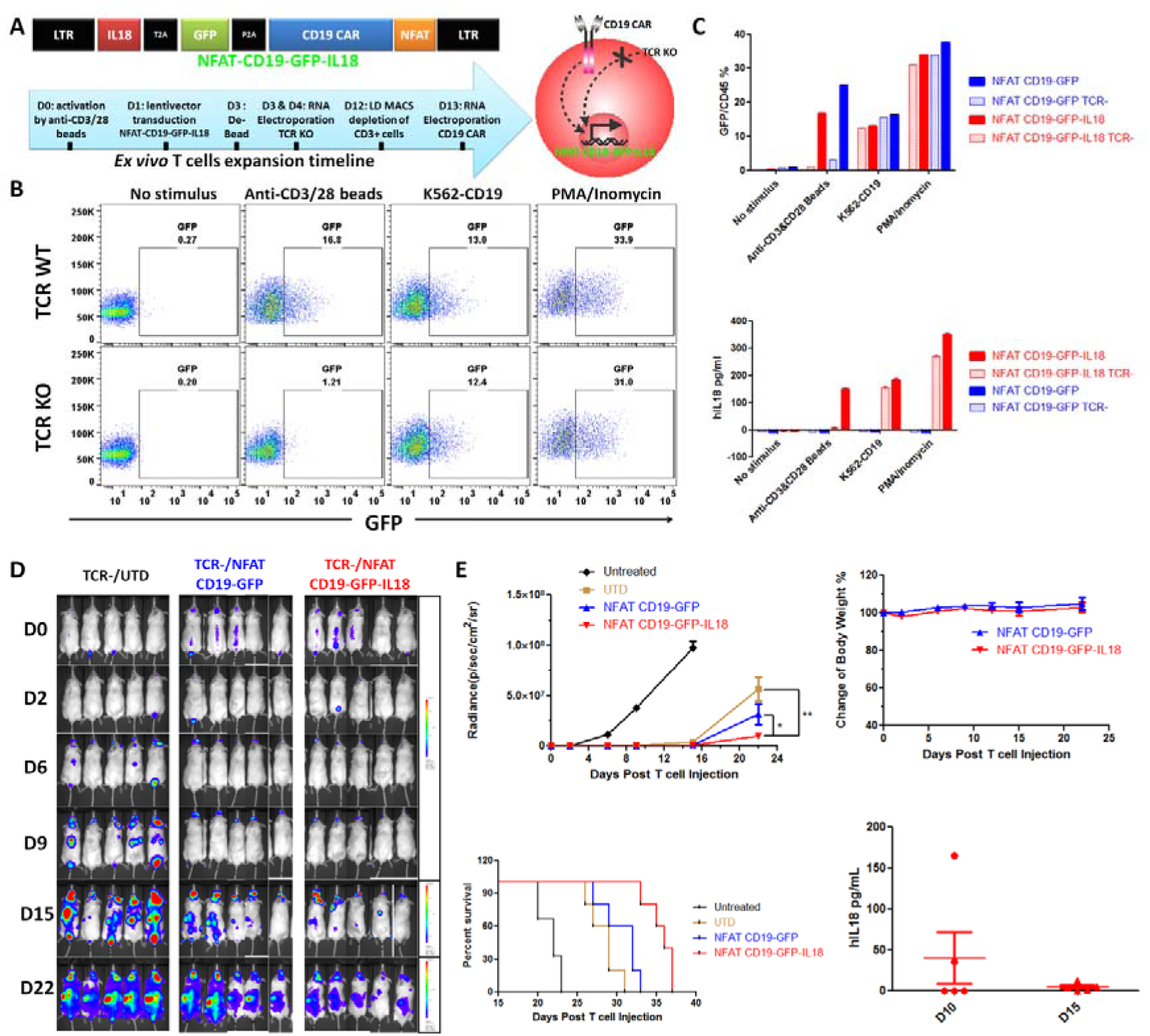
An inducible system for IL18 expression in CART cells. **(A)**Upper panel: schematic of lentiviral construct in which the NFAT promoter drives expression of CD19-GFP-IL18. Lower panel: the *ex vivo* timeline to generate TCR-deficient T cells transduced with NFAT-CD19-GFP-IL18 and electroporated with mRNA CD19 CAR (designated as TCR-/NFAT-CD19-GFP-IL18). **(B)** TCR-/NFAT-CD19-GFP-IL18 CART cells were stimulated with anti-CD3/28 beads, K562-CD19, or PMA/ionomycin, after which CD45GFP+ cells were estimated by flowcytometry. **(C)** Summary of GFP+/CD45+ percentages (upper panel) and IL18 concentrations (lower panel) under different conditions as indicated. **(D)** NSG mice (n=5) were inoculated with 1×10^6^ Nalm6-CBG/GFP cells, and treated 1 week later with TCR-deficient untransduced, NFAT-CD19-GFP, or NFAT-CD19-GFP-IL18 CART (all bearing mRNA encoded CD19 CAR). Tumor progression was monitored by live animal imaging. **(E)** Tumor growth kinetics (Student's t-test: **p=0.0057, *p=0.0359), survival rate after T cells infusion, change of body weight, and IL18 concentrations at D10 and D15 following T cell injection. All data with error bars are presented as mean±SEM.

We tested the inducible NFAT-CD19-GFP-IL18 TCR-deficient T cells expanded as described in the timeline shown in Figure 6A. TCR-/NFAT-CD19-GFP-IL18 T cells were restimulated under different conditions, and analyzed for GFP expression and IL18 production. The results showed that irradiated K562-CD19 aAPC and PMA/ionomycin treatment induced similar levels of GFP expression and IL18 synthesis in TCR WT and TCR KO NFAT-CD19-GFP-IL18 T cells, whereas anti-CD3/28 beads only activated GFP expression and IL18 synthesis in TCR WT NFAT-CD19-GFP-IL18 T cells (Figure 6B,C).This observation demonstrated that the inducible system, which we deemed **A**ntigen **P**ropelled **A**ctivation of **C**ytokine **H**elper **E**nsemble (APACHE), operates as intended. In NSG mice bearing Nalm6 xenograft tumors, APACHE-armored CD19 CART cells suppressed tumor progression more effectively, and resulted in better survival, than TCR-/NFAT CD19-GFP CART cells and TCR-/untransduced (UTD) T cells expressing mRNA encoded CD19 CAR (Figure 6D,E). APACHE-armored CD19 CART cells did not induce significant loss of body weight and only low levels of systemic IL18 were detected at day 10 and day 15 following T cell injection (Figure 6E), suggesting that this inducible system could minimize the potential risk of continuous systemic IL18 expression.

## Discussion

Except for B cell tumors treated with CD19 CAR T cells, a major limitation of current CAR T cell approaches has been insufficient expansion and persistence of the infused T cells, as exemplified by trials in ovarian and brain cancer (Brown et al., 2015; Kershaw et al., 2006), and reviewed in (Fesnak et al., 2016; Gilham et al., 2012; Jensen and Riddell, 2015). IL12, a cytokine related to IL18, has shown promise in augmenting antitumor effects and T cell expansion (Chmielewski et al., 2014; Pegram et al., 2012). There may be some advantages to the use of IL18, including that it is a single chain cytokine in contrast to IL12, which is heterodimeric. In addition, IL18 has been tested as a single agent in clinical trials and shown to have moderate antitumor effects and an acceptable safety profile (Robertson et al., 2006). In contrast, IL12 has been shown to have unacceptable toxicity in previous trials (Cohen, 1995).

IL18 can regulate both innate and adaptive immune responses through its effects on natural killer (NK) cells, monocytes, dendritic cells, T cells, and B cells (Dinarello et al., 2013). The most important biologic effect of IL18 is that it acts synergistically with other pro-inflammatory cytokines to promote interferon-Γ production by NK cells, T cells, and possibly other cell types. Due to the pleiotropic functions of IL18, tissue-accumulated IL18 has been shown to impact both the tumor stroma and the immune response, by sustaining T cell cytolytic activity, recruiting and activating innate immune cells, and re-programming stroma associated immune suppressor cells (Chmielewski et al., 2014). The rationale for our studies is that the accumulation of IL18 as a transgenic pro-inflammatory cytokine is intended to recruit a second wave of immune cells to the tumor microenvironment to mediate antitumor effects towards cancer cells that are resistant to the direct effects of the CAR T cells.

Many studies have shown that efficacy of adoptive cell therapy with tumor targeted T cells is enhanced by host conditioning using radiation or lymphodepleting chemotherapy. Intensive chemotherapy or radiation given to enhance adoptive cell therapy has been reported to have severe toxicity in some patients (Goff et al., 2016). Given the pronounced augmentation of CAR T cell engraftment that we observed with T cells secreting IL18, it is possible that this may decrease the need for prior host conditioning. In preclinical models with mouse CAR T cells engineered to secrete IL-12, the need for lymphodepletion was obviated (Pegram et al., 2012).

One concern with CAR T cells armored with IL18 is the potential for systemic inflammation (Car et al., 1999). To mitigate this possibility several strategies will be employed in clinical trials. First, CAR-IL18 T cells will be given in low doses where the calculated maximal levels of IL18 could not reach the levels shown to be safe in humans in previous trials (Robertson et al., 2006). Secondly, if cytokine levels become a concern, they can be reduced by incorporation of various kill switch strategies (Straathof et al., 2003). Finally, a more elegant approach was developed here where the IL18 transgene was not constitutively expressed, but rather, was only induced and secreted upon encountering with the surrogate tumor antigen. Presumably this would limit IL18 secretion to the tumor microenvironment, and limit systemic exposure.

A limitation of our *in vivo* studies is that they were conducted in an immunodeficient NSG mouse model. A strength of this model is that it enabled testing of the APACHE CAR T cells in a robust tumor model. However, the tumor microenvironment is lacking in this model since the tumor stroma is composed largely of mouse cells. In humans and in mice, the biologic activity of IL18 is balanced by the presence of a high affinity, naturally occurring IL18 binding protein (IL18BP). This binding occurs with a high affinity and has been reported to occur interspecies between mouse and human IL18 (Aizawa et al., 1999). It is possible that the toxicity observed in the mice manifested by systemic inflammation, weight loss and death in some cases, was exaggerated or underreported based on the relative balance of mouse IL18BP and the transgenic IL18 secreted by the APACHE T cells.

In summary, we demonstrate that CAR T cells can be engineered to secrete IL18 that supports *in vivo* engraftment and persistence of the CAR T cells. Our observations demonstrate that IL18 has a previously unappreciated form of costimulation on human CD4+ T cells, and therefore these IL18 expressing CD4 T cells can be categorized as synthetic T helper cell producing IL18 (sTH18). These findings may have therapeutic implications because the CAR IL18 T cells can support the proliferation of CD8 T cells, and because increasing evidence points to the importance of adoptively transferred CD4+ T cells in antitumor effects (Hunder et al., 2008; Hung et al., 1998). Finally, CAR T cells with inducible IL18 may have a role in immune-surveillance and have a therapeutic potential for inducing antigenic spread. Before translation of this technology into clinical setting, further investigation is underway to address the IL18-CART toxicity and biology in syngeneic models.

## Materials and Methods

### Mice and cell lines

NOD-SCID-G chain-/- (NSG) were purchased from Jackson Laboratories, and all animal procedures were performed in animal facility at the University of Pennsylvania in accordance with Federal and Institutional Animal Care and Use Committee requirements. Wild-type parental AsPC1, K562 cells and Nalm-6 cells (purchased from American Type Culture Collection) were engineered to express CBG-T2A-GFP as a reporter gene and trace of luciferase activity *in vitro* and *in vivo.* K562 with CBG-T2A-GFP cells were further engineered to express human CD19 (K562-CD19) or human mesothelin (K562-CD19).

### Molecular cloning

The parental vector pTRPE SS1BBz contains scFV recognizing human mesothelin, 4-1BB and CD3z intracellular signaling domains. The sequence encoding mature human IL18 (Uniprot Q14116) were synthesized from IDT and cloned into pTRPE SS1BBz along with a T2A (Kim et al., 2011) linker (pTRPE SS1BBz-T2A-hIL18). GFP and Humanized CAR CD19BBz were cloned to replace SS1BBz to generate pTRPE CD19BBz-T2A-hIL18 and pTRPE GFP-T2A-hIL18. GFP was cloned to replace SS1BBz to generate pTRPE GFP. GFP was also cloned to replace hIL18 to generate pTRPE SS1BBz-T2A-GFP and pTRPE CD19BBz-T2A-GFP. In order to generate the lentivector construct pTRPE NFAT CD19-GFP-IL18, the parental vector pTRPE SS1BBz was depleted of the fragment consisting EF1A promoter and SS1 BBz, and inserted with a reverse sequence of synthesized fragment consisting of an NFAT promoter (Zhang et al., 2015) followed by GFP with a P2A (Kim et al., 2011) linker. The resulting pTRPE NFAT GFP vector was modified by replacing GFP with CD19BBz, which was further modified by inserting a fragment of GFP or GFP-T2A-IL18 downstream of P2A site to generate pTRPE NFAT CD19-GFP or CD19-GFP-IL18.

### Primary T cell expansion

Primary donor T cells were transduced with lentivector encoding CAR or GFP with or without hIL18 and expanded as previously reported (Milone et al., 2009). In brief, primary donor T cells were activated with anti-CD3/CD28 Dynabeads (LifeTechnologies) at D0, and transduced with lentivector at D1. Fresh medium was added at D3 and the activated T cells were de-beaded at D5 and maintained at 7e5/ml until rested T cells were frozen in liquid nitrogen.

### CRISPR/Cas9 gene knockout

Primary donor T cells were transduced with lentivector encoding CAR or GFP with or without hIL18 and engineered with CRISPR/Cas9 system as previously reported (Cong et al., 2013; Jinek et al., 2012; Mali et al., 2013; Ren et al., 2016). In brief, primary donor T cells were activated with anti-CD3/CD28 Dynabeads (LifeTechnologies) at D0, and transduced with lentivector at D1. The activated T cells were de-beaded at D3 and electroporated with sgRNA and Cas9 RNA at D3 and D4. Then the T cells were maintained at 7e5/ml until rested T cells were frozen in liquid nitrogen.

TCR-deficient T cells were achieved with sgRNA targeting GGAGAATGACGAGTGGACCC within TCRB region. For the IL18R knockout we designed 7 sgRNA targeting IL18RA (Cong et al., 2013; Jinek et al., 2012; Mali et al., 2013), and utilized RNA electroporation to induce CRISPR/Cas9 mediated gene editing following the procedure of generating TCR KO universal CART (Ren et al., 2015). The screening of IL18RA KO showed that a the A4 sgRNA produced the most efficient gene knockout (Figure 4-figure supplement 1A), and sequencing of the PCR product spanning the gene edited site confirmed mutations in the DNA sequence (GTACAAAAGCAGTGGATCAC) targeted by the A4 sgRNA (Figure 4-figure supplement 1B).

## Abbreviation

TCR: T Cell Receptor
CAR: Chimeric Antigen Receptor
IL18-CART: IL18 producing CART
CD19-IL18 CART: IL18 producing CD19 CART
SS1-IL18 CART: IL18 producing SS1 CART
TIL: Tumor Infiltrating Lymphocytes
GVHD: Graft-Versus-Host Disease
CRISPR: Clustered regularly interspaced short palindromic repeats
Cas9: CRISPR associated protein 9
IL18R: Knockout
KO: IL18 Receptor
NFAT: Nuclear Factor of Activated T-cells
aAPC: artificial Antigen Presenting Cells

## Contributions

CHJ and BH proposed the concept of IL18 expressing CART for cancer immunotherapy. BH designed all, and performed most, of the experiments. JR, BH, and YZ executed the knockout of TCR and IL18R by CRISPR/Cas9. BH and YL conducted live animal imaging, Trucount analyses and mouse necropsy. JS provided scientific resources, such as parental cloning vectors and research animals. BH, CHJ, BK and RMY wrote and edited the manuscript.

## Acknowledgements

This work was supported by a US National Institutes of Health (NIH) grant to C.H.J and Y.Z (2R01CA120409), by a sponsored research grant from Novartis, and National Institutes of Health National Cancer Institute grants T32 CA009140 (B.H.). This arrangement is managed in accordance with the University of Pennsylvania’s Conflict of Interest Policy. The authors are in compliance with this policy.

Throughout the study, we really appreciate excellent scientific coordination by Regina Young, and passionate discussion and great help from Chien-Ting Lin and Avery Posey in the June lab. We also appreciate all the help from other members in the June Lab, including Decheng Song, Joseph Fraietta, Shannon McGettigan, Sonia Guedan, Kathleen Haines, Tong Da, Keisuke Watanabe, Mauro Castellarin, Christopher Kloss and Michael Klichinsky. We are really grateful to the analysis of whole mouse pathology by Elizabeth Buza and her team from Comparative Pathology Core at UPENN, and the luminex analysis by Patricia Tsao and Yinan Lu from Human Immunology Core at UPENN, and the IHC staining by Daniel Martinez from Pathology Core Laboratory of Children’s Hospital of Philadelphia Research Institute.

B.H., C.H.J. and Y.Z. have financial interests due to a pending patent application. Conflicts of interest are managed in accordance with University of Pennsylvania policy and oversight. The other authors declare that they have no competing interests.

The following figure supplements are available for figure 1:

**Figure supplement 1.** SS1-IL18 CART cells displayed enhanced cytotoxicity.

**Figure supplement 2.** IL18 expanded T cells are polyclonal through phenotyping of TCRB variants.

**Figure supplement 3.** CD19-IL18 CART cells mediate dramatic T cell expansion and lethal GVHD in xenograft model.

The following figure supplements are available for figure 2:

**Figure supplement 1.** Cytokine profile of IL18 and anti-CD3 beads activated T cells.

**Figure supplement 2.** Memory phenotype of T cells treated with recombinant IL18 and anti-CD3 beads.

**Figure supplement 3.** Memory phenotype of anti-CD3/IL18 restimulated T cells and aAPC restimulated IL18-CART cells.

The following figure supplements are available for figure 3:

**Figure supplement 1.** Characterization of SS1-IL18 CART cells during extended proliferation.

**Figure supplement 2.** Cytokine production by IL18-CART cells stimulated with aAPC.

**Figure supplement 3.** IL18RA expression and memory phenotype of IL18 expanded T cells in xenograft model.

The following figure supplements are available for figure 4:

**Figure supplement 1.** CRISPR/Cas9 targeted knockout of IL18RA expression.

The following figure supplements are available for figure 5:

**Figure supplement 1.** Characterization of relapsing Nalm6 cells and re-expanded CD19-IL18 TCR-CART cells.

